# Methanogenic Ethanol Production from Acetate

**DOI:** 10.64898/2026.07.07.736952

**Authors:** Ruchira Mitra, Hyeon-Ji Hwang, Yunjeong Choi, Ingmar H. Riedel-Kruse, Thomas K. Wood

**Affiliations:** Department of Chemical Engineering, Pennsylvania State University, University Park, Pennsylvania, 16802-4400, USA; Department of Molecular and Cellular Biology, University of Arizona, Tucson, AZ, 85721-0020, USA; Departments of Applied Mathematics, Biomedical Engineering, and Physics, University of Arizona, Tucson, AZ, 85721-0020, USA

**Keywords:** Methanogen, ethanol, acetate, aceticlastic methanogenesis, reverse methanogenesis

## Abstract

Biological ethanol production is important for the circular carbon economy and makes up 73% of the U.S. biological fuels market. Previously, we produced ethanol by reversing methanogenesis and capturing methane by cloning methyl-coenzyme M reductase (Mcr) from an unculturable population of anaerobic methanotrophic archaea (ANME-1); this process was predicated on the generation of the intermediate acetate and its conversion by the methanogenic host to ethanol. Moreover, methanogens are generally thought to be detrimental for converting acetate to ethanol and are usually intentionally inhibited. Here, we demonstrate for the first time the conversion of acetate as the sole carbon and energy source to both ethanol and methane by the methanogen *Methanosarcina acetivorans* C2A, resulting in 40% of the metabolized acetate becoming ethanol. Moreover, we found there is 14-fold more ethanol produced from acetate by *M. acetivorans* C2A compared to that from methane and 5-fold more ethanol compared to *M. acetivorans* C2A producing ANME-1 Mcr using methane. As expected, we found growth on methanol results primarily in methane generation and low levels of ethanol. Therefore, acetate may be readily converted by the methanogen *M. acetivorans* to ethanol at high yields, along with methane production.

## INTRODUCTION

As the global demand for sustainable biofuels is escalating, ethanol has emerged as a cornerstone for transitioning towards a low-carbon economy (Vacharanukrauh et al., 2025). Ethanol is widely used in transportation sector (Balat & Balat, 2009), pharmaceutical (Fine-Shamir & Dahan, 2024), personal care and cosmetics (Celikoglu et al., 2025), food and beverage (Herdiana, 2025), and green chemical industries (Vacharanukrauh et al., 2025). The conventional first-generation ethanol production relies on edible feedstock such as corn, and sugarcane (Prasad et al., 2019). The United States, accounting for ∼56% (Althuri & Venkata Mohan, 2022) of the global ethanol production, uses ∼40% (Ramsey et al., 2023) of its corn harvest to produce ethanol. This has sparked significant debates regarding agricultural land-use competition, deforestation, and food insecurities (Kanso et al., 2025) thereby necessitating more sustainable approaches that leverages circular carbon cycles (Vacharanukrauh et al., 2025).

Methane is a potent greenhouse gas, 28-times more detrimental than carbon dioxide on a per-molecule-basis in trapping atmospheric heat (Wood et al., 2023). Most of the current methane emission (∼60%) is anthropogenic, primarily from enteric fermentation, livestock manure management, rice cultivation, landfills and wastewater management, biomass burning, oil and gas drilling sites, and coal mining (Tucci & Rosenzweig, 2024). Methane being the most reduced form of carbon, it also represents an important energy source for the global economy (Guerrero-Cruz et al., 2021) and, thus capturing methane via microbial oxidation is a highly coveted research objective in recent years. Methanotrophs aerobically oxidize ∼30 Tg yr^−1^of methane to methanol (Tucci & Rosenzweig, 2024) whereas, a consortium of anaerobic methanotrophic archaea (ANME-1) inhabiting the marine sediments anaerobically oxidizes up to an order of magnitude more methane (∼300 Tg/yr) with sulfate, nitrate, or metal ions as electron acceptors (Knittel & Boetius, 2009), thereby preventing a massive quantity of methane from entering the atmosphere from the ocean.

The enzyme catalyzing methane capture is methyl-coenzyme M reductase (Mcr) (Shima et al., 2012). A major drawback of directly utilizing these archaea for anaerobic oxidation of methane (AOM) is their inability to grow as pure culture, due to their long lag phase (∼ 60 years) and doubling time (∼ seven months) (Wood et al., 2023). A significant advance in AOM was achieved, by our research group, when the *mcrBGA* genes encoding the ANME-1 Mcr were cloned into a genetically-tractable methanogen *Methanosarcina acetivorans* (Soo et al., 2016). The heterologously-expressed ANME-1 Mcr (pES1-MAT*mcr3*) enabled the host to capture methane and carbon dioxide and convert them to acetate via reverse methanogenesis, allowing for the first time anaerobic growth on methane by a pure culture (Soo et al., 2016); this breakthrough was predicated on the discovery that Fe^+3^ could be used as the terminal electron acceptor (Soo et al., 2016). Using Fe^+3^ as the electron acceptor, additional insights into the biochemistry for growth on methane by *M. acetivorans* by reversing methanogenesis have been determined (Yan et al., 2018), and growth on methane by reversing methanogenesis with Fe^+3^ as the electron acceptor and the formation of acetate by *M. acetivorans* has been corroborated (Yan et al., 2023).

We further engineered this recombinant *M. acetivorans* host by cloning 3-hydroxybutyryl-CoA dehydrogenase (Hbd) from *Clostridium acetobutylicum* to produce *L*-lactate from the captured methane (McAnulty et al., 2017b). We have also constructed a synthetic consortium utilizing the recombinant *M. acetivorans* host with cloned ANME-1 Mcr, methane-acclimated sludge, and *Geobacter sulfurreducens* to create the first microbial fuel cell converting captured methane directly into electricity (McAnulty et al., 2017a; Yamasaki et al., 2018); notably, this system produces electricity from methane as well as any biofuel cell (Yamasaki et al., 2018). Recently, using biochar as the electron acceptor, methanogenesis was reversed with *M. acetivorans* to synthesize polyhydroxybutyrate (Li et al., 2025). Hence, *M. acetivorans* producing Mcr is a proven chassis for green chemistry.

Recently, we discovered that the acetate produced by the same host, *M. acetivorans* with cloned ANME-1 Mcr, from the captured methane and carbon dioxide (dissolved as bicarbonate), is further converted to ethanol by utilizing its native aldehyde ferredoxin oxidoreductase (AOR) and alcohol dehydrogenase (Adh) with ferric iron as the electron acceptor (Mitra et al., 2026; Soo et al., 2016):

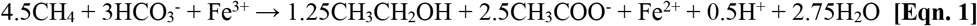

This production of ethanol from growth on methane using Fe^+3^ as the electron acceptor by reversing methanogenesis with *M. acetivorans* was recently corroborated (Yan et al., 2026). Moreover, we increased ethanol production via the reversal of methanogenesis with cloned ANME-1 Mcr by silencing the chromosomal copy of Mcr that primarily produces methane via RNAi (Hwang et al., 2026a) as well as by performing the first directed evolution of ANME-1 Mcr (Hwang et al., 2026b).

Ethanol may be produced through acetate reduction during the fermentation of organic waste using anaerobic sludge as inoculum and hydrogen as the electron donor (Steinbusch et al., 2008); however, methanogenesis is intentionally inhibited to enhance ethanol production (Steinbusch et al., 2009; Steinbusch et al., 2008). Moreover, growth on acetate by *Methanosarcina* spp. is considered to produce primarily methane, not ethanol (Ferry, 2020). The carbonyl group of the acetate is oxidized and the released electrons are transferred to the endogenous methyl group producing methane (Prakash et al., 2019). The stoichiometry for acetate reduction to methane via aceticlastic methanogenesis is (Ferry, 2020):

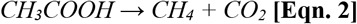

Surprisingly, we demonstrate here that acetate is readily converted to both ethanol and methane, rather than exclusively to methane, by the methanogen *M. acetivorans*. We also propose that reverse methanogenesis of the acetate-derived methane to produce acetate provides the electrons required for driving the acetate reduction to ethanol. To the best of our knowledge, this is the first report of the conversion of acetate to both ethanol and methane by a methanogen.

## MATERIALS AND METHODS

### Archaeal strain and culture conditions

*M. acetivorans* C2A (DSMZ Collection #2834, henceforth “C2A”) and the engineered strain *M. acetivorans* C2A/pES1-MAT*mcr3* (Amp^R^, Pur^R^, R6K *ori*, C2A *ori*, P_mcr_ANME-1_::*mcrBGA*_ANME-1_, henceforth “C2A/pES1-MAT*mcr3*”) (Soo et al., 2016) were routinely grown anaerobically at 39 ^°^C in high salt (HS) medium buffered with 50 mM 1,4-piperazinediethanesulfonic acid (PIPES) (Soo et al., 2016) and 48 mM sodium carbonate supplemented with 2.5 g/L yeast extract (HSYE) with methanol (0.5% v/v) as the carbon source and Na_2_S·9H_2_O (0.025% w/v) as the reducing agent; puromycin (2 µg/mL) was used to maintain plasmid pES1-MAT*mcr3* in C2A (Soo et al., 2016). The pH of the medium was 6.5. HSYE medium (100 mL) was inoculated with a thawed 1.8 mL aliquot of each glycerol stock in an anaerobic chamber, the bottles were sealed, and cultures were incubated at 200 rpm for 7 to 10 days to a turbidity at 600 nm of ∼ 0.5.

For the conversion of acetate (100 mM) to ethanol experiments, methanol-grown cells were concentrated to a turbidity of 3.5 by centrifugation at 5,000 rpm for 20 min. The cell pellet was washed three times to remove residual methanol with HS medium containing 48 mM sodium carbonate, 0.025% w/v Na_2_S·9H_2_O. Washed cell pellets were then resuspended in 12 mL of HS medium containing 48 mM sodium carbonate, and 0.025% w/v Na_2_S·9H_2_O in 40 mL glass vials. The crimp-sealed vials were then incubated at 250 rpm. Similarly, for the conversion of methanol to ethanol experiments, methanol-grown cells were concentration to turbidity of 3.5, washed, and resuspended in 12 mL of HS medium containing 0.5% v/v methanol, and 0.025% w/v Na_2_S·9H_2_O in 40 mL glass vials.

For the growth on methane experiments, freshly prepared and filter-sterilized (0.22 μm) FeCl_3_ (10 mM) was added to the culture as the electron acceptor, and the glass vials were crimped. The liquid phase was then sparged with 99% of methane at 70 mL/min for 5 min using both inlet and outlet needles to allow continuous gas exchange and prevent pressure buildup. The headspace (28 mL) was fully replaced with methane, and the pressure was maintained at atmospheric pressure. The crimp-sealed vials were then incubated inverted at 250 rpm to prevent gas loss.

For the growth on acetate experiments, thawed 1.8 mL aliquot of C2A glycerol stock was inoculated in 100 mL of HS or HSYE medium containing acetate (50 mM), 48 mM sodium carbonate, and Na_2_S·9H_2_O (0.025% w/v) in an anaerobic chamber. The bottles were sealed, and incubated at 200 rpm, and growth was monitored by measuring the OD_600_.

### Electroporation of *M. acetivorans*

Plasmid pES1-MAT*mcr3* was transformed into *M. acetivorans* by electroporation under anaerobic conditions (Mitra et al., 2026). Cells (10 mL) grown in HSYE medium (with 0.5% v/v methanol and 0.025% w/v Na_2_S·9H_2_O) to a turbidity at 600 nm of 1.0 were harvested by centrifugation at 8000 rpm for 15 min and washed twice in same volume of cold (0°C) 0.85 M sucrose. Washed cells, resuspended in 100 μL of cold 0.85 M sucrose, mixed with 5 μg of plasmid DNA was electroporated (1.25 kV, 200 Ω, and 25 μF) under anaerobic conditions in a cold electroporation cuvette using a BioRad Gene Pulser apparatus (model number 165-2098). Cells mixed with 900 μL HSYE medium (with 0.5% v/v methanol and 0.025% w/v Na_2_S·9H_2_O) was then transferred to 3 to 5 mL HSYE medium (with 0.5% v/v methanol and 0.025% w/v Na_2_S·9H_2_O) and incubated for 24 h for recovery. The cells were then harvested by centrifugation, resuspended in 10 mL HSYE medium (with 0.5% v/v methanol, 0.025% w/v Na_2_S·9H_2_O and 2 μg/mL puromycin) and incubated for 5 to 7 days to obtain the transformed cells. Transformation was validated by polymerase chain reaction (PCR) of the puromycin resistance gene present in the plasmid backbone using the forward primer 5’-CAGCAACAGATGGAAGGCC-3’ and the reverse primer 5’-TCGTAGAAGGGGAGGTTGC-3’.

### Total protein estimation

Total protein was quantified using the Bradford assay (Bio-Rad Laboratories, Hercules, CA, USA) (Mitra et al., 2026). Samples (100 µL) were harvested by centrifugation at 6,000 g for 5 min, and the supernatant and cell pellet were separated. Cell pellets were resuspended in distilled water to a final volume of 100 µL, vortexed for 30 sec, and heated at 90°C for 60 min for cell lysis. Supernatants and the cell lysate suspensions were transferred to a 96-well plate, mixed with 200 µL of 1× Bradford reagent, and the absorbance was recorded at 595 nm.

### Methane, methanol, ethanol, and acetate assays via gas chromatography

All gas chromatography (GC) analyses were performed on Agilent 6890N gas chromatograph using nitrogen as the carrier gas (Mitra et al., 2026). Methane was analyzed using Carboxen-1000 packed column (4600 × 2.1 mm, Supelco catalog no. 12390-U) and thermal conductivity detector (TCD). Injections were 30 µL from the headspace. The temperatures of front inlet and detector were maintained at 150°C and 240°C, respectively. The column temperature was initially set at 185°C, then increased to 220°C at a rate of 16°C/min and held at 230°C for 6 min. Nitrogen served as both carrier and reference gas.

Methanol and ethanol were assayed using 0.1% AT-1000 packed column (6′, 1/8″ O.D., 80/100 mesh on Graphpac-GC support; Alltech) and flame ionization detector (FID). The temperatures of front inlet and detector were maintained at 150°C and 210°C, respectively. The column temperature was initially set at 50°C for 1 min, then increased to 220°C at a rate of 50°C/min and held for 2 min. The supernatant (2 µL) of samples harvested by centrifugation at 8000 g for 8 min was injected. Concentrations were calculated from a calibration curve prepared using standard methanol or ethanol solutions in the same medium composition as used in the experiments.

Acetate was assayed using a Stabilwax-DA capillary column (30 m × 0.53 mm i.d., 0.25 µm film; Restek) with FID detection. The front inlet and detector were set to 250°C. The oven program was 70°C hold, increase to 100°C at 40°C/min, and then increase to 200°C at 100°C/min. Supernatant of the sample was acidified with 1% v/v formic acid and then 2 µL was injected. The calibration curve was prepared with standard solution of sodium acetate in the same medium composition as used in the experiments.

### Statistical analysis

Statistical analysis was performed using GraphPad Prism 10. Data are represented as the mean ± standard deviation (SD) of at least three independent cultures. Student’s T-test was performed to evaluate the difference between data sets, with probability values (p) < 0.05 considered significant.

## RESULTS

### Rationale

Previously, we showed that C2A/pES1-MAT*mcr3*, engineered to capture methane and carbon dioxide via ANME-1 Mcr, produces the biofuel precursor, acetate (Soo et al., 2016), which is further converted to ethanol, probably via its native aldehyde ferredoxin oxidoreductase and alcohol dehydrogenase (Mitra et al., 2026). We, thus, hypothesized that the wild type C2A, if grown directly on acetate, might be able to utilize this acetate to produce ethanol using its native aldehyde ferredoxin oxidoreductase and alcohol dehydrogenase. Therefore, we cultivated C2A on acetate for six days and assayed acetate consumption, methane production, and ethanol production from that acetate. Subsequently, we compared it to methanol-grown C2A as well as methane-grown-C2A/pES1-MAT*mcr3*.

### Ethanol production from acetate

With acetate as the sole carbon source at an initial input of 1110 μmol, we found ethanol was produced by C2A at high concentrations after six days (140 ± 30 μmol). Normalized ethanol, based on the number of cells present as indicated by the total protein level, was 37 ± 8 μmol/mg **(Fig. 1A)**. The acetate consumption was 340 ± 90 μmol (**Fig. 1B**). To investigate if methane was generated, we measured the headspace methane level in the closed vials and found that methane production was 280 ± 1 μmol (**Fig. 1B**). Hence, our hypothesis was confirmed, and acetate consumed by the C2A is readily converted to both ethanol as well as methane.

**Fig 1.**
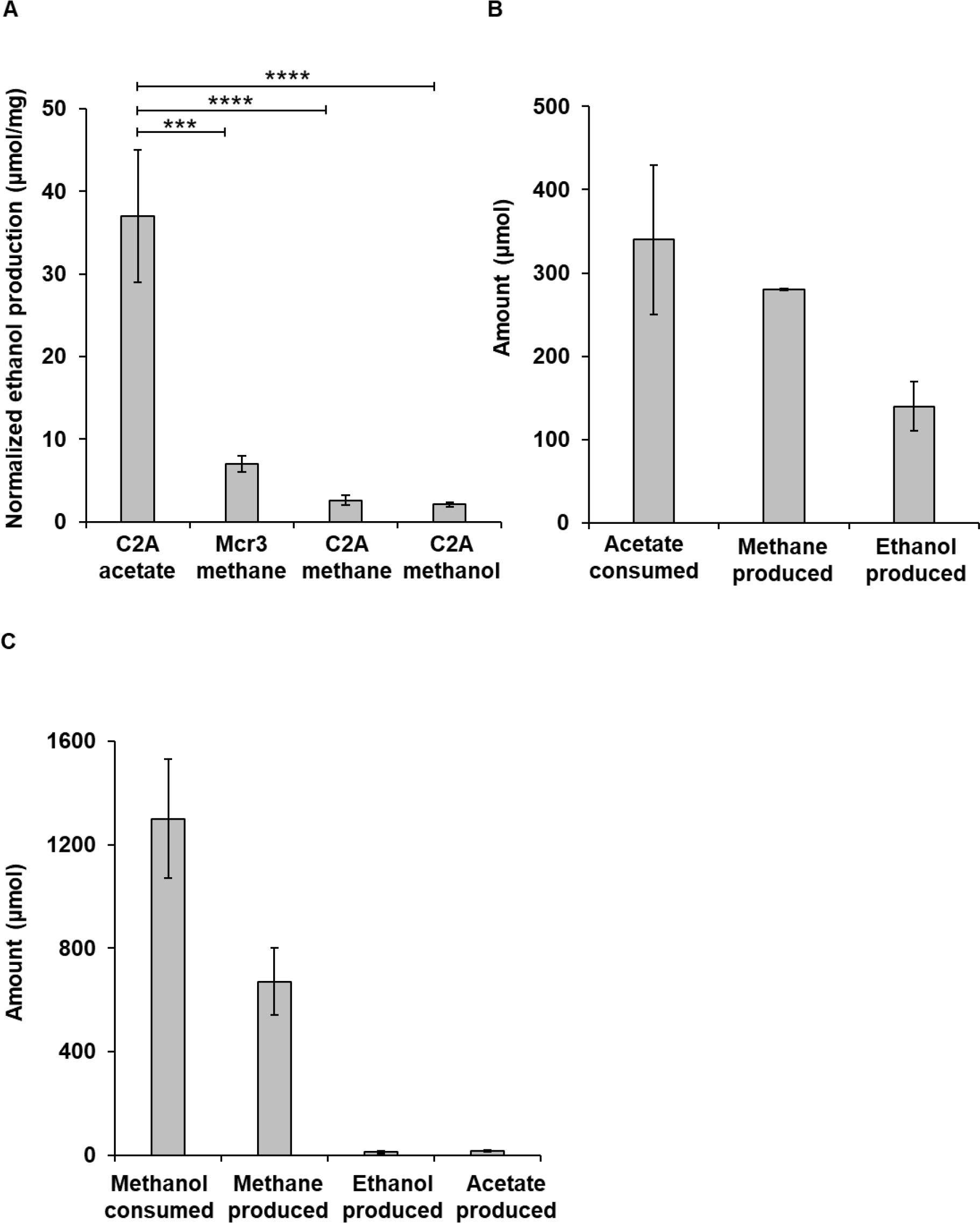
Ethanol production by *M. acetivorans* from acetate. **(A)** Normalized ethanol produced (i) from acetate by *M. acetivorans* C2A (C2A acetate), (ii) from methane by *M. acetivorans* C2A/pES1-MAT*mcr3* (Mcr3 methane), (iii) from methane by *M. acetivorans* C2A (C2A methane), and (iv) from methanol by *M. acetivorans* C2A (C2A methanol). C2A is wild-type *M. acetivorans* producing Mcr that primarily generates methane during methanogenesis, and Mcr3 is *M. acetivorans* C2A/pES1-MAT*mcr3* producing ANME-1 Mcr that captures methane during reverse methanogenesis. Ethanol values are normalized by total protein after six days. Bars and error bars are the mean and standard deviation of at least three independent cultures with C2A on acetate five independent cultures. *** p < 0.005, **** p < 0.0005. **(B)** Acetate consumption, methane production, and ethanol production by C2A after six days of incubation on 0.1 M acetate-containing HS medium. **(C)** Methanol consumption, methane production, ethanol production, and acetate production by C2A after six days of incubation on 0.5% v/v methanol-containing HS medium.

### Stoichiometric analysis supports acetate to ethanol and methane conversion

We estimate (**Table 1**) the stoichiometry for acetate conversion to ethanol by *M. acetivorans* as:

**Table 1.**
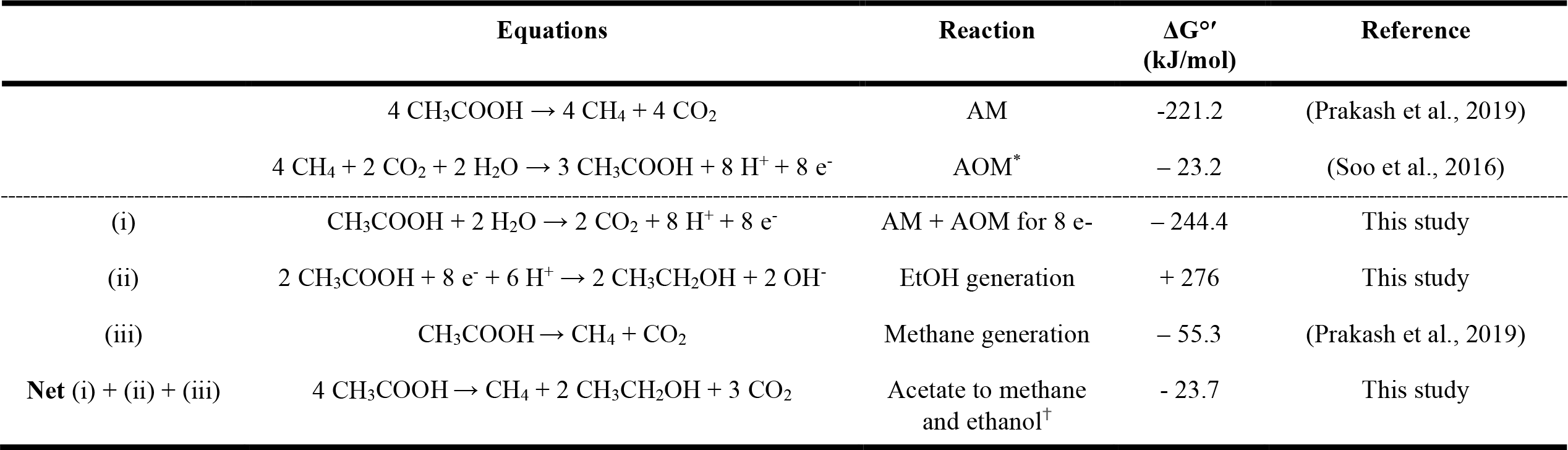
Stoichiometry and thermodynamics for the acetate-to-methane and acetate to ethanol pathway in *M. acetivorans*. Standard Gibbs free energies (ΔG°′) are expressed under standard biochemical conditions (pH 7, 25°C, 1 atm). AM, aceticlastic methanogenesis; AOM, anaerobic oxidation of methane by reverse methanogenesis. ^*^For the AOM reaction, the Gibbs free energy (Soo et al., 2016) was transformed to ΔG°′ at pH 7 using ΔG°′ = ΔG° + RT ln([H^+^]^8^). ^†^For the conversion of acetate to methane and ethanol, the net stoichiometric equation and Gibbs free energy are obtained by summing the reactions (i) + (ii) + (iii).

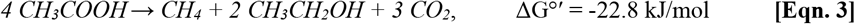

which includes using one mol of acetate to generate eight electrons that are required to reduce two mol of acetate to two mol of ethanol via aceticlastic methanogenesis combined with anaerobic oxidation of methane (**Fig. 2**). Experimentally, from 340 ± 90 μmol of acetate, 140 ± 30 μmol of ethanol and 280 ± 1 μmol of methane was produced (**Fig. 1B**). Clearly, ∼40% of the total carbon was recovered in the ethanol, and the methane production accounts for another ∼40% of the total carbon. Since, for every two moles of ethanol, three moles of carbon dioxide are released as byproduct (**Eqn. 3**), the remaining input carbon may have converted to carbon dioxide. Based on the predicted carbon dioxide production from **Eqn. 3**, the carbon recovery was calculated to 113%.

**Fig 2.**
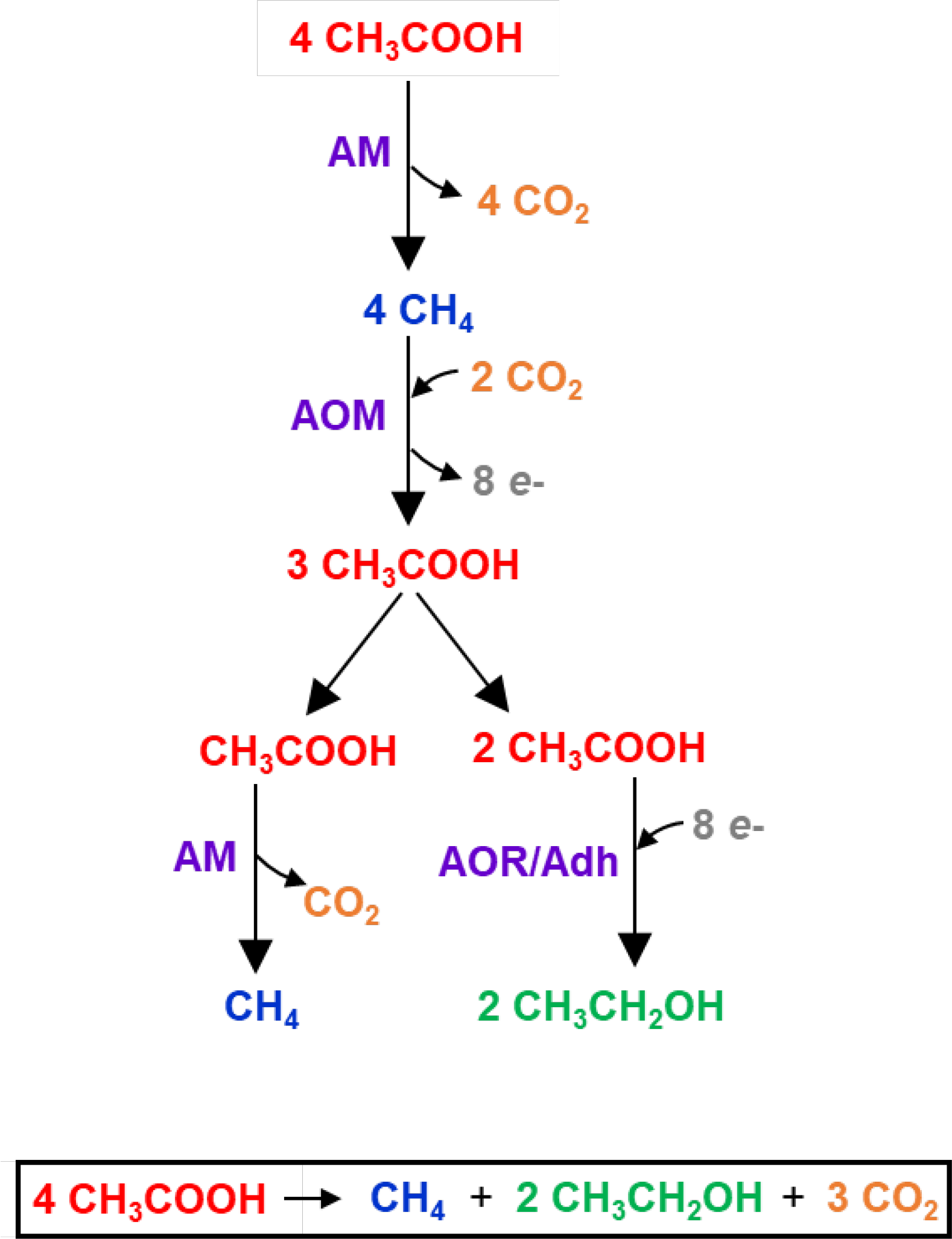
Schematic for the conversion of acetate to methane, carbon dioxide, and ethanol by *M. acetivorans* C2A. Simplified representation of the electron flow for the conversion of acetate to methane and carbon dioxide via aceticlastic methanogenesis (AM), methane conversion to acetate via anaerobic oxidation of methane (AOM) by reverse methanogenesis, and acetate conversion to ethanol via native aldehyde ferredoxin oxidoreductase (AOR) and alcohol dehydrogenase (Adh). Metabolites: CH_3_COOH, acetic acid; CO_2_, carbon dioxide, CH_4_, methane; and CH_3_CH_2_OH, ethanol.

We also checked for possible contamination using microscopy and found that the cultures we used are pure C2A and C2A/pES1-MAT*mcr3* (**Fig. 3A**). Furthermore, we also checked for growth of C2A on acetate and confirmed that C2A uses acetate for growth only in the presence of yeast extract, as originally published (Sowers et al., 1984) (**Fig. 3B**). Since we did not use yeast extract for the six-day conversion of acetate to ethanol, our results are related to conversion, not growth.

**Fig 3.**
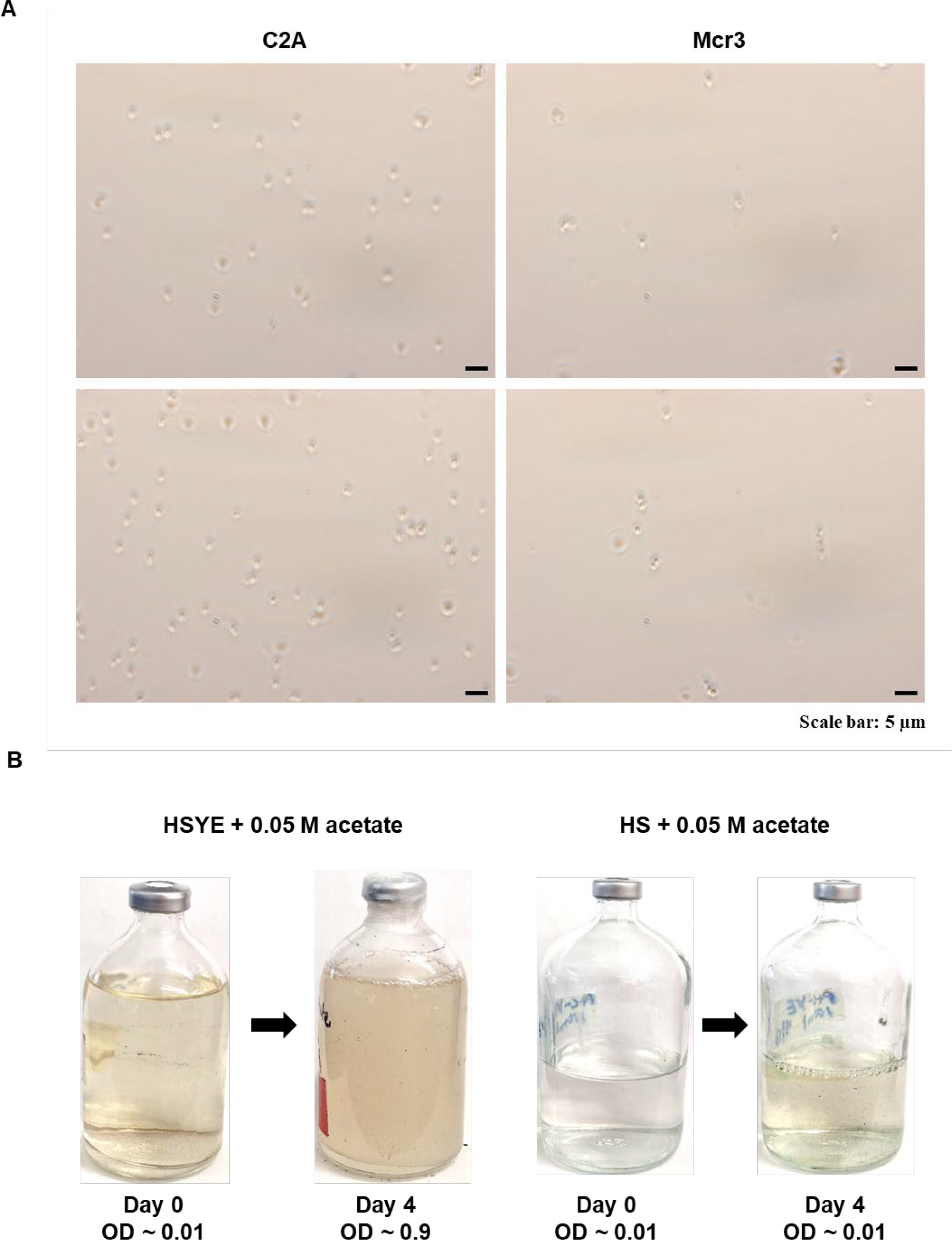
Microscopic observation of *M. acetivorans* C2A and *M. acetivorans* C2A/pES1-MAT*mcr3* under methanol-grown conditions and growth of *M. acetivorans* C2A on acetate. **(A)** Representative bright-field micrographs (100× magnification) of *M. acetivorans* C2A (C2A) and C2A/pES1-MAT*mcr3* (Mcr3) cultures grown with methanol as the sole carbon and energy source. Images were acquired using a Zeiss Axio Scope A1 microscope. Although *M. acetivorans* C2A/pES1-MAT*mcr3* cells appeared slightly smaller than *M. acetivorans* C2A, no substantial differences in cell morphology were observed between the two strains under methanol-grown conditions. Scale bars = 5 μm. **(B)** Growth of *M. acetivorans* C2A on acetate (0.05 M). HSYE represents high-salt medium containing 2.5 g/L of yeast extract, and HS represents high-salt medium without yeast extract.

### Thermodynamic analysis suggests coupling of reverse methanogenesis and acetate reduction to ethanol

Thermodynamic analysis indicates that the acetate conversion to ethanol and methane is exergonic (ΔG°′ = -22.8 kJ/mol), and thus thermodynamically feasible **(Table 1, Eqn. 3)**. We further reasoned that the reducing power required for acetate reduction to ethanol is provided through the oxidation of methane (derived from acetate via aceticlastic methanogenesis) to acetate via reverse methanogenesis **(Fig. 2)**. Thermodynamic analysis supports this coupling mechanism. Aceticlastic methanogenesis to convert acetate to methane and carbon dioxide is exergonic (ΔG°′ = -221.2 kJ/mol) and reverse methanogenesis to oxidize methane back into acetate is also exergonic (ΔG°′ = -23.2 kJ/mol) **(Table 1)**. These two exergonic reactions enabled the reduction of acetate to ethanol.

### C2A yields low amounts of ethanol from methanol

Methanol is the most-preferred carbon source for *Methanosarcina* species and is routinely used to culture C2A. Hence, we analyzed if C2A produces ethanol using methanol as the carbon source. Low levels (11 ± 4 μmol) of ethanol were produced using methanol as the sole carbon source (**Fig. 1C**). Normalized ethanol was 2.1 ± 0.3 μmol/mg **(Fig. 1)**. We found that entire methanol (1300 ± 230 μmol) was consumed by C2A after six days (**Fig. 1C**). Corroborating this methanol consumption, the headspace methane in the closed vials after six days was 670 ± 130 μmol (**Fig. 1C**), which was 2.4 -fold higher than that produced from acetate. According to the stoichiometric equation (Yin et al., 2019), the calculated Gibbs free energy shows that the conversion of methanol to methane is highly exergonic:

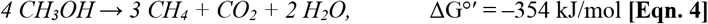

The experimental methane produced correlated well with the theoretical methane yield (975 μmol). Considering the predicted carbon dioxide production from **Eqn. 4**, the carbon recovery was calculated to 69%. To investigate if the methane produced was converted to the intermediate acetate, we measured the acetate levels and found that low amounts (16 ± 3 μmol) were produced. Hence, as expected, the methanol consumed by the C2A was primarily converted to methane via highly exergonic methylotrophic methanogenesis but only low levels of acetate and ethanol was produced.

### C2A produces 5-fold higher ethanol from acetate than C2A/pES1-MATmcr3

To compare the ethanol produced C2A from acetate to engineered C2A/pES1-MAT*mcr3* from methane, we cultivated C2A/pES1-MAT*mcr3*, at the same cell density, on methane for six days. The ethanol produced by C2A/pES1-MAT*mcr3* was 36 ± 8 μmol (cf., 140 ± 30 μmol by acetate-grown C2A). Consistent with previously-reported normalized ethanol production by high-cell densities of C2A/pES1-MAT*mcr3* (8.1 ± 0.8 μmol/mg) (Mitra et al., 2026), the normalized ethanol by C2A/pES1-MAT*mcr3* was 7 ± 1 µmol/mg (vs. 37 ± 8 μmol/mg by C2A contacted with acetate) **(Fig. 1)**. Hence, ethanol produced by C2A from acetate was 5-fold better than the ethanol produced by growing C2A/pES1-MAT*mcr3* on methane at similar turbidity. Furthermore, the normalized ethanol by the methane-grown C2A was 2.6 ± 0.6 µmol/mg **(Fig. 1)**, thus, ethanol by C2A contacted with acetate was 14-fold better than the methane-grown C2A and substantial amounts of ethanol may be recovered from acetate by wild-type C2A.

## DISCUSSION

Our results demonstrate ethanol may be generated from acetate, along with the previously-reported methane (Ferry, 2020), by the methanogen *M. acetivorans* when contacted with acetate as the sole carbon source. We found that the ethanol produced by acetate-grown wild type *M. acetivorans* is five-fold higher than the ethanol produced by the methane-grown engineered *M. acetivorans* with cloned ANME-1 Mcr and 14-fold better than the methane-grown wild-type *M. acetivorans* (**Fig. 1**). We also found that low levels of acetate and ethanol are produced by methanol-grown wild type *M. acetivorans* with methane as the primary product.

We recently showed that *M. acetivorans* with cloned ANME-1 Mcr is capable of producing ethanol by capturing methane and carbon dioxide, with acetate as the intermediate (Mitra et al., 2026). We also reported that the wild-type *M. acetivorans* expressing the native Mcr is capable of producing ethanol by capturing low amounts of methane and carbon dioxide, via formation of the intermediate acetate (Mitra et al., 2026). Since acetate is the key precursor for ethanol production, it is reasonable that when *M. acetivorans* is contacted directly with acetate in this report (rather than generating it from methane), significantly higher levels of ethanol are achieved compared to the methane-grown *M. acetivorans*. Unlike the original report that acetate is primarily converted to methane (Ferry, 2020), our experimental results and carbon recovery analysis indicate that 40% of the acetate-derived carbon was converted to methane and another 40% was converted to ethanol. Our discovery here of the production of ethanol from acetate is due to the high cell density used along with the utilization of our previous discovery that C2A has active enzymes for converting acetate to ethanol (Mitra et al., 2026).

Maintaining redox balance is critical for the growth of methanogens. We propose that the reducing power necessary for acetate reduction to ethanol is supplied via reverse methanogenesis of methane back to acetate; basically, acetate is split in two with the oxidized half becoming CO_2_ and the other, reduced half becoming methane that undergoes reverse methanogenesis to generate the electrons needed to reduce acetate to ethanol (**Fig. 2**). Specifically, owing to the ability of *M. acetivorans* to perform both methanogenesis and reverse methanogenesis **(Fig 2)**, we suggest that 4 mol of acetate from the acetate pool is initially converted to 4 mol of methane and 4 mol of carbon dioxide via aceticlastic methanogenesis **(Fig. 2)**. These 4 mol of methane and 2 mol of carbon dioxide undergo reverse methanogenesis producing 3 mol of acetate, generating 8 electrons to reduce the 2 mol of acetate to ethanol via the native AOR/Adh **(Fig. 2)**. The remaining 1 mol of acetate is converted to 1 mol of methane and 1 mol of carbon dioxide. Thus, a net 1 mol of acetate generate 8 electrons with 2 mol of carbon dioxide **(Table 1** and **Fig. 2)**. Therefore, overall, 4 mol of acetate generates 2 mol of ethanol, 1 mol of methane, and 3 mol of carbon dioxide. Hence, this work demonstrates that acetate may be converted to both ethanol and methane by methanogens for biotechnological applications. In addition, this work has implications for understanding the role of methanogenic organisms in environments like cold seeps where acetate is generated from methane by anaerobic methanotrophic archaea (Yang et al., 2020) and our work suggests the methanogens present may convert some of this acetate to ethanol.

## CREDIT AUTHORSHIP CONTRIBUTION STATEMENT

**Ruchira Mitra:** Investigation, Methodology, Data curation, Writing - original draft, Writing - review & editing. **Hyeon-Ji Hwang:** Investigation, Data curation. **Yunjeong Choi:** Data curation. **Ingmar H. Riedel-Kruse:** Fund acquisition, Writing - review & editing. **Thomas K. Wood:** Supervision, Conceptualization, Project administration, Fund acquisition, Writing - original draft, Writing - review & editing.

## DECLARATION OF COMPETING INTERESTS

The authors declare no competing interests.

## ACKNOWLEDGEMENTS

We thank J. Cuello, R. Sierra Alvarez, and members of the Riedel-Kruse Lab for stimulating discussions.

## FUNDING STATEMENT

This work was supported by funds provided by the NSF-FMRG grant 2229070.

## DATA AVAILABILITY

All data are contained within the manuscript.

## Notes

### Competing Interest Statement

The authors have declared no competing interest.

### Summary of Updates

The thermodynamic analysis has been updated and proof has been added that the cultures are pure.

